# Biological sex and BMI influence the longitudinal evolution of adolescent and young adult MRI-visible perivascular spaces

**DOI:** 10.1101/2024.08.17.608337

**Authors:** Erin A. Yamamoto, Seiji Koike, Caitlyn Wong, Laura E. Dennis, Madison N. Luther, Avery Scatena, Seva Khambadkone, Jeffrey J. Iliff, Miranda M. Lim, Swati R. Levendovszky, Jonathan E. Elliott, Giuseppe Barisano, Eva M. Müller-Oehring, Angelica M. Morales, Fiona C. Baker, Bonnie J. Nagel, Juan Piantino

## Abstract

**Background and Purpose:** An association recently emerged between magnetic resonance imaging (MRI)-visible perivascular spaces (MV-PVS) with intracerebral solute clearance and neuroinflammation, in adults. However, it is unknown how MV-PVS change throughout adolescence and what factors influence MV-PVS volume and morphology. This study assesses the temporal evolution of MV-PVS volume in adolescents and young adults, and secondarily evaluates the relationship between MV-PVS, age, sex, and body mass index (BMI).

**Materials and Methods:** This analysis included a 783 participant cohort from the longitudinal multicenter National Consortium on Alcohol and Neurodevelopment in Adolescence study that involved up to 6 imaging visits spanning 5 years. Healthy adolescents aged 12-21 years at study entry with at least two MRI scans were included. The primary outcome was mean MV-PVS volume (mm^3^/white matter cm^3^).

**Results:** On average, males had greater MV-PVS volume at all ages compared to females. A linear mixed-effect model for MV-PVS volume was performed. Mean BMI and increases in a person’s BMI were associated with increases in MV-PVS volume over time. In females only, changes in BMI correlated with MV-PVS volume. One unit increase in BMI above a person’s average BMI was associated with a 0.021 mm^3^/cm^3^ increase in MV-PVS volume (p<0.001).

**Conclusion:** This longitudinal study showed sex differences in MV-PVS features during adolescence and young adulthood. Importantly, we report that increases in BMI from a person’s mean BMI are associated with increases in MV-PVS volume in females only. These findings suggest a potential link between MV-PVS, sex, and BMI that warrants future study.

## INTRODUCTION

Magnetic resonance imaging (MRI)-visible perivascular spaces (MV-PVS) are observed throughout the lifespan, including in healthy youth.[1,2] In adults, the MV-PVS has drawn significant attention as a marker of cerebral small vessel disease and age-related neurodegenerative processes,[3] and have been associated with three key neurologic processes– immune surveillance,[4,5] CSF and interstitial fluid exchange,[6–9] and intracerebral solute clearance.[9,10] The role of MV-PVSs in maintaining homeostasis in the young and developing brain remains understudied.

The young adolescent brain undergoes significant structural changes that follow mapped developmental trajectories. By age 6, the brain achieves 95% of its peak size,[11] but total brain volume is greatest at 10.5 years and 14.5 years in girls and boys, respectively.[12] Gray matter and white matter (WM) mature differently, where WM increases throughout childhood and gray matter follows an inverted U-shaped trajectory ultimately decreasing in volume in adolescence.[11] The development of MV-PVS in youth was modeled in one cross-sectional study,[1] but no longitudinal studies have been conducted. Understanding the longitudinal progression of MV-PVS development may provide insight into the presence of MV-PVS in healthy adolescents and young adults.

Sex differences have been observed in brain development, neurologic disease, and also in MV-PVS burden.[2,12–15] Males have greater brain volume than females,[11,12,14] and boys were shown to have greater MV-PVS burden in children.[1,2] Males and females also have different susceptibilities to neurologic disease processes.[16,17] Some conditions that predominantly affect young females, such as multiple sclerosis and neuromyelitis optica,[18] migraines,[19] and idiopathic intracranial hypertension[20,21] are associated with increased MV-PVS burden relative to controls.[22–25] Clinically, obesity has been proposed as a mediator of disease severity in these female-dominant conditions, and was also associated with elevated intracranial pressure and chronic inflammation.[26,27] BMI, a surrogate measure of obesity, was associated with increased MV-PVS burden and was the most significant variable correlated with MV-PVS volume in a young adult population.[1,15,28] The interactions between sex, BMI, and MV-PVS burden in young individuals have not been well-studied.

In a recent cross-sectional study, we observed lower MV-PVS volume in females than male adolescents,[2] a finding replicated in another cross-sectional study.[1] While novel, the interpretation and practical utility of these early findings were limited by 1) cross-sectional design that does not account for individual differences over time, 2) use of distinct research cohorts,[1] 3) no assessment of MV-PVS within the basal ganglia,[2] and 4) lack of BMI as a covariate in sex regression models.

Using a longitudinal neuroimaging database of healthy adolescents and young adults, we aimed to 1) establish the temporal evolution of MV-PVS for a given individual and 2) determine the relationship between sex, BMI, and MV-PVS in youth. We hypothesized that males and females would significantly differ in MV-PVS volume and the trajectory of MV-PVS burden over time. Moreover, given that MV-PVSs have been associated with both female-predominant neurologic conditions and BMI, we hypothesized that changes in BMI would have a differential effect on MV-PVS burden in males and females.

## MATERIALS AND METHODS

### Study Design

Subjects were enrolled in the National Consortium on Alcohol and Neurodevelopment in Adolescence, a multisite, cohort-sequential, longitudinal study.[29] Annual study visits and neuroimaging occurred at study entry and up to 7 years of follow-up at 5 study sites.

### Standard Protocol Approvals and Patient Consents

All study subjects provided informed assent/consent. The local institutional review board approved the use of de-identified data from the National Consortium on Alcohol and Neurodevelopment in Adolescence.

### Study Participants

Healthy volunteers aged 12-21 years at study entry were eligible to participate. Cross-sectional analysis from participants at a single site were previously reported.[2] Exclusion criteria included the following: inability to understand English; residence >50 miles from an assessment site; MRI contraindications; current use of medications that affect brain function or blood flow (e.g. antidepressants, stimulants); history of serious medical problems that could affect MRI (e.g. diabetes, recurrent migraine, traumatic brain injury with loss of consciousness for > 30 minutes); noncorrectable vision or hearing impairments; mother who drank >2 alcoholic drinks in a week or used nicotine >10 times per week, marijuana >2 times per week, or other drugs during pregnancy; prematurity (<30 weeks’ gestation); low birth weight or other perinatal complications requiring intervention; current diagnosis of Axis I psychiatric disorder; substance dependence; learning disorder; other developmental disorder requiring specialized education.[29] Participants with a single MRI timepoint were excluded from the current analysis (Supplemental Figure 1). Clinical and demographic variables were collected at study entry (hereon referred to as "baseline"). BMI was calculated from self-reported height and weight at each visit.

### MR Imaging Acquisition and Preprocessing

T1- and T2-weighted 3D images were collected on 4 scanner models: 3T General Electric Discovery MR750 and SIGNA Creator, and 3T Siemens Tim Trio and Prisma Fit. Images were collected in the sagittal plane on scanners from 2 manufacturers – 3T General Electric Discovery MR750 (3 sites) and SIGNA Creator (1 site), and 3T Siemens Trio Tim (2 sites) and Prisma Fit scanners (2 sites). Sites using the GE scanners used an 8-channel head coil. 3D T1-weighted sequences were acquired by Inversion Recovery-Spoiled Gradient Recalled (IR-SPGR) echo sequence (TR/TE = 5.904/1.932 ms, TI = 400 ms, flip angle = 11°, matrix = 256 x 256, FOV = 24 cm, voxel dimensions = 1.2 x 0.9375 x 0.9375 mm, 146 slices) and 3D Sagittal CUBE T2 sequence (TR/TE = 2500/99.646 ms, ETL=100, matrix = 512 x 512, FOV = 240 x 240 mm, voxel dimensions = 1.2 x 0.4688 x 0.4688 mm, 160 slices). At sites using the Siemens scanner, a 12-channel head coil was used to acquire a 3D T1-weighted gradient echo sequence (TR/TE = 1900/2.92 ms, TI = 900 ms, flip angle = 9°, matrix = 256 x 256, FOV = 240 mm, voxel dimensions = 1.2 x 0.9375 x 0.9375mm, 160 slices) and a 3D T2-weighted turbo spin-echo with variable excitation pulse sequence (TR/TE = 3200/404 ms, matrix = 256 x 256, FOV = 240 x 240 mm, voxel dimensions = 1.2 x 0.9375 x 0.9375mm, 160 slices). Three individual human “phantoms” traveled between and were repeatedly scanned at each of the five sites. Imaging preprocessing steps were described previously.[30] Briefly, T1 and T2 weighted images were corrected for gradient nonlinearity, readout, and bias field.[31] FreeSurfer longitudinal stream (Version 7.4) was used to generate WM, cortical gray matter, basal ganglia (BG), and ventricular CSF segmentations.

### PVS Segmentation and Quantification

The MV-PVS quantification method used here has been described previously.[30] Briefly, T1- and T2-weighted images were combined to enhance the MV-PVS signal contrast and denoised with adaptive non-local mean filtering to remove high-frequency spatial noise.[32]

MV-PVS were segmented using a multiscale vessel enhancement filtering approach with default parameters.[32,33] Quantification of MV-PVS volume, count, diameter, and solidity was performed on the WM and BG masks. MV-PVS total volume (mm^3^/cm^3^) and total count (number/cm^3^) were expressed as a ratio to mask volume to account for differences in brain size.[28] Diameter was expressed as the mean MV-PVS diameter (mm) within a given mask of interest. Solidity is a measure of MV-PVS morphology, ranging from 0 to 1, where 1 indicates a perfectly linear MV-PVS and less than 1 suggests curvature to the MV-PVS.[30]

### Statistical Analysis

Statistical analyses were performed using STATA/MP 18 (StataCorp). The cohort’s baseline demographic and MV-PVS characteristics by sex were analyzed with descriptive statistics. Continuous variables are reported as mean [standard deviation]. MV-PVS characteristics were assessed by age, BMI, and sex. The age by age variable represents a quadratic effect of age, to account for a potentially non-linear relationship between age and MV-PVS burden. Differences in baseline characteristics between sexes were analyzed by independent samples t-test. Locally weighted scatter plot smoothing curves were derived from the subject data (3649 observations: 1849 female, 1800 male) to visualize MV-PVS trajectories over the cohort’s range of ages and BMIs for each MV-PVS parameter.

Based on prior studies, MV-PVS volume was selected as the parameter of interest.[1,2] Age, sex, BMI and scanner model were included as covariates in a linear mixed-effects model. Age and BMI were centered on the cohort mean across all timepoints, 18-years-old and 23 kg/m^2^, respectively. BMI was evaluated by person-mean BMI and person-occasion BMI, representing an individual’s average BMI across all timepoints and an individual’s deviation from their mean BMI at each timepoint, respectively. The effect of sex, person-mean BMI, and person-occasion BMI deviation were examined before exploring potential interactions. All models were estimated using residual maximum likelihood. The significance of fixed effects were determined by Wald test p-values (two-sided), while likelihood ratio tests (one-sided) determined the significance of additional random effects. The threshold for statistical significance was set to ⍺=0.05.

### Data Availability Statement

Data are available from the National Consortium on Alcohol and Neurodevelopment in Adolescence study team on request. The code used to analyze these data is available on request.

## RESULTS

### Baseline study characteristics

The final sample consisted of 783 individuals (Supplemental Figure 1). The average age was 16.2 [2.6] years at study enrollment, and 50.3% of the cohort was female (Table 1). The mean BMI at enrollment was 22.2 [4.3] kg/m^2^. There were no sex differences in age or BMI. MV-PVS were identified by previously published algorithm (Figure 1).[30] Males had significantly greater mean MV-PVS volume (7.26 [1.23] mm^3^/WM cm^3^ vs. 6.65 [1.26] mm^3^/WM cm^3^, p<0.001), mean MV-PVS count per WM cm^3^ (1.59 [0.29] vs. 1.52 [0.27], p<0.001), and mean MV-PVS diameter (1.79 [0.09] mm vs. 1.77 [0.08] mm, p=0.002) than females. There were no sex differences in MV-PVS solidity (Table 1).

**Figure 1.**
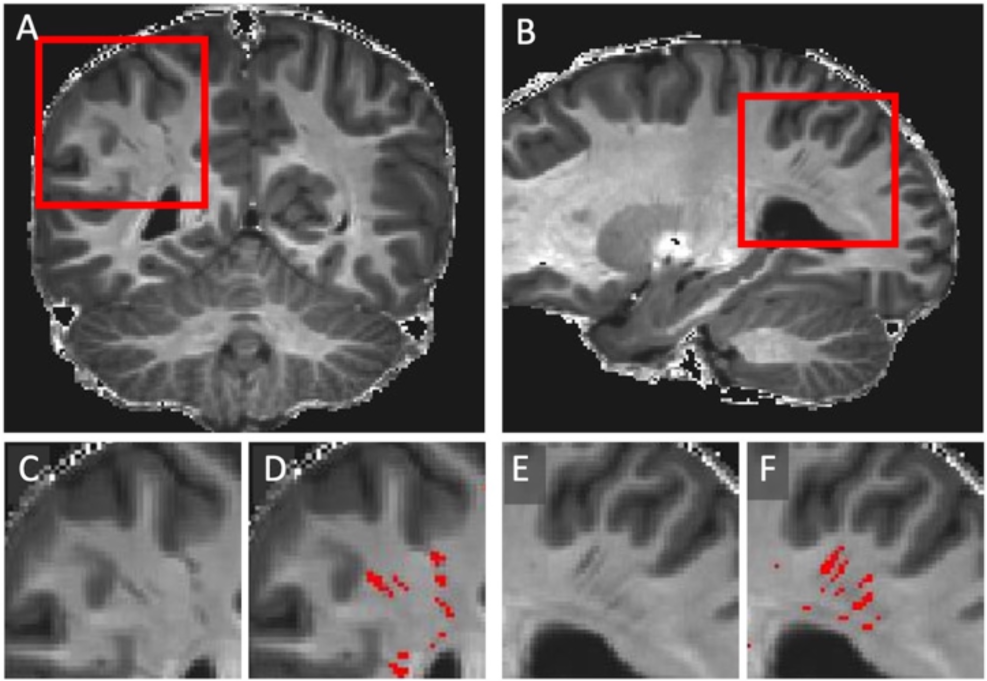
Example of the PVS detection algorithm. T1- and T2-weighted images were filtered and combined to enhance the visibility of PVS structures, as shown in the coronal (A, C-D) and sagittal planes (B, E-F) demonstrating MV-PVS detection. The red inset boxes (A and B) mark PVS regions of interest. These regions are magnified (C and E, respectively), then displayed with a PVS mask overlay in red (D and F, respectively).

**Table 1.**
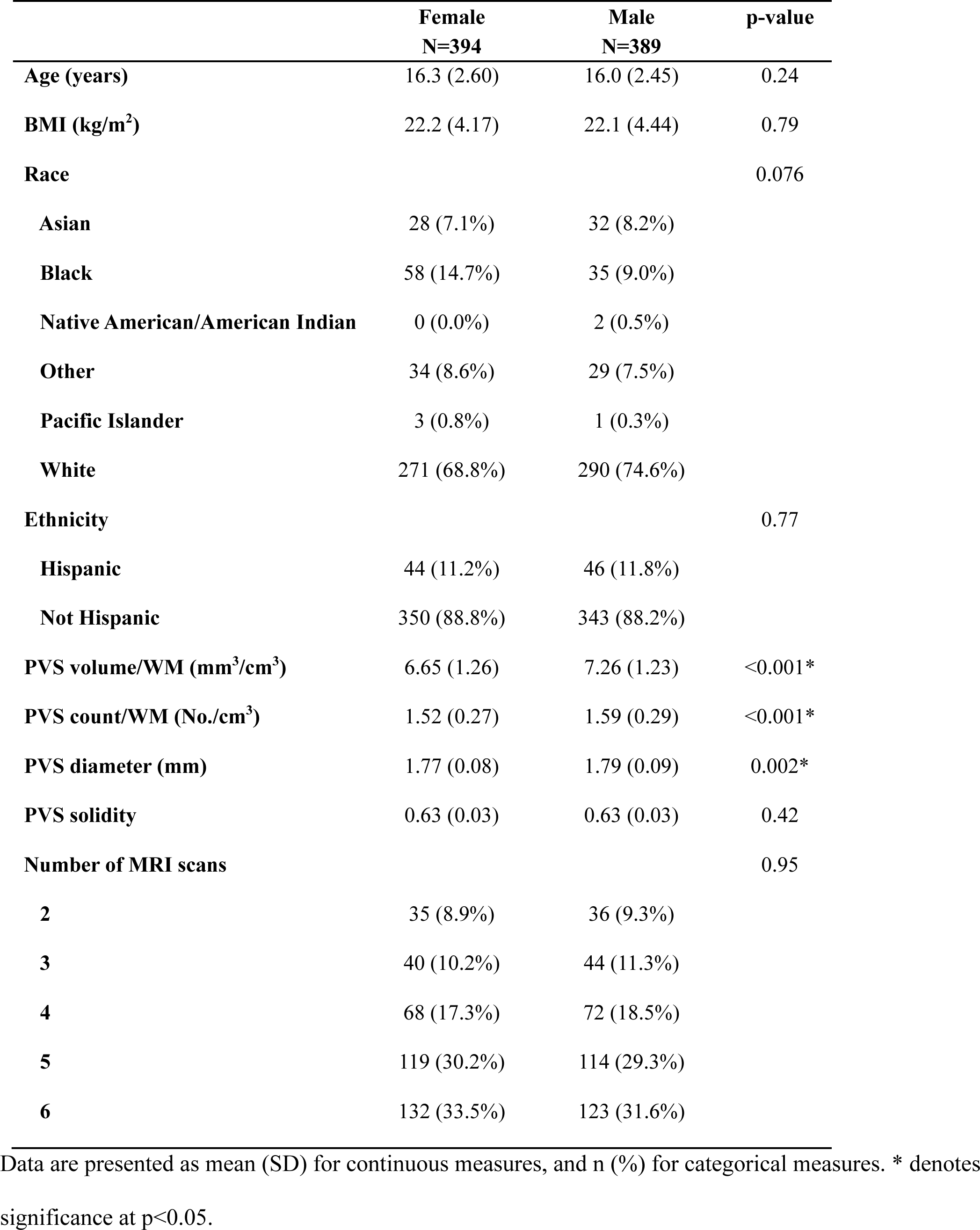
Baseline cohort characteristics.

|  | **Female**<br>**N=394** | **Male**<br>**N=389** | **p-value** |
| --- | --- | --- | --- |
| **Age (years)** | 16.3 (2.60) | 16.0 (2.45) | 0.24 |
| **BMI (kg/m<sup>2</sup>)** | 22.2 (4.17) | 22.1 (4.44) | 0.79 |
| **Race** |  |  | 0.076 |
| **Asian** | 28 (7.1%) | 32 (8.2%) |  |
| **Black** | 58 (14.7%) | 35 (9.0%) |  |
| **Native American/American Indian** | 0 (0.0%) | 2 (0.5%) |  |
| **Other** | 34 (8.6%) | 29 (7.5%) |  |
| **Pacific Islander** | 3 (0.8%) | 1 (0.3%) |  |
| **White** | 271 (68.8%) | 290 (74.6%) |  |
| **Ethnicity** |  |  | 0.77 |
| **Hispanic** | 44 (11.2%) | 46 (11.8%) |  |
| **Not Hispanic** | 350 (88.8%) | 343 (88.2%) |  |
| **PVS volume/WM (mm<sup>3</sup>/cm<sup>3</sup>)** | 6.65 (1.26) | 7.26 (1.23) | <0.001* |
| **PVS count/WM (No./cm<sup>3</sup>)** | 1.52 (0.27) | 1.59 (0.29) | <0.001* |
| **PVS diameter (mm)** | 1.77 (0.08) | 1.79 (0.09) | 0.002* |
| **PVS solidity** | 0.63 (0.03) | 0.63 (0.03) | 0.42 |
| **Number of MRI scans** |  |  | 0.95 |
| **2** | 35 (8.9%) | 36 (9.3%) |  |
| **3** | 40 (10.2%) | 44 (11.3%) |  |
| **4** | 68 (17.3%) | 72 (18.5%) |  |
| **5** | 119 (30.2%) | 114 (29.3%) |  |
| **6** | 132 (33.5%) | 123 (31.6%) |  |

Data are presented as mean (SD) for continuous measures, and n (%) for categorical measures. \* denotes significance at p<0.05.

### Cohort trends of MV-PVS development by age and sex in subcortical white matter and basal ganglia

We first sought to describe the pattern of observed longitudinal MV-PVS progression from individual subject observations (Figure 2-3). In WM (Figure 2), males had greater MV-PVS volume compared to females at all ages (Figure 2A). By cross-sectional analyses, there were no statistical differences in MV-PVS volume between males and females at ages 12 years and younger. Males also had greater MV-PVS counts with a divergence between sexes at approximately age 17 years (Figure 2B). Mean MV-PVS diameter increased with age. Males had larger mean MV-PVS diameter than females in adolescence but were comparable to females moving into early adulthood (Figure 2C). Cross-sectional analysis showed no difference in mean diameter between sexes at 20 years. Notably, increasing MV-PVS volume and count are not secondary to decreasing WM volume (Supplemental Figure 2). In contrast to the increase in MV-PVS observed in WM, BG MV-PVS volume and count decreased with age (Figure 3A-B). BG MV-PVS diameter increased with age and was not significantly different between males and females (Figure 3C). Solidity remained constant with age (Figure 3D).

**Figure 2.**
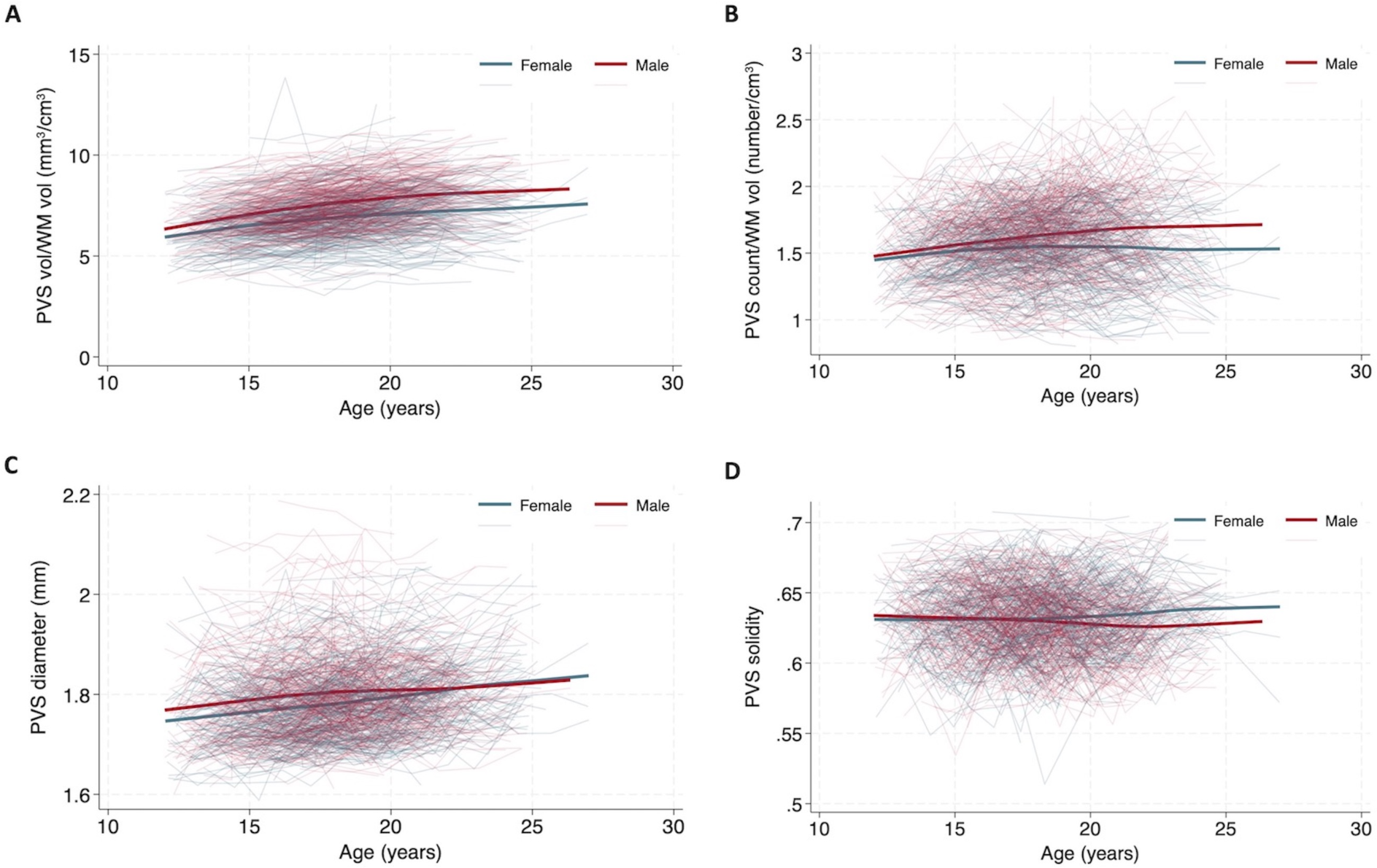
Relationship between white matter MV-PVS burden and age. Changes in WM MV-PVS parameters A) volume B) count, C) diameter, D) solidity over time are represented for each individual by spaghetti plot. Locally weighted scatter plot smoothing curves derived from 3649 observations (1849 female-blue, 1800 male-red) are also shown. WM vol = white matter volume.

**Figure 3.**
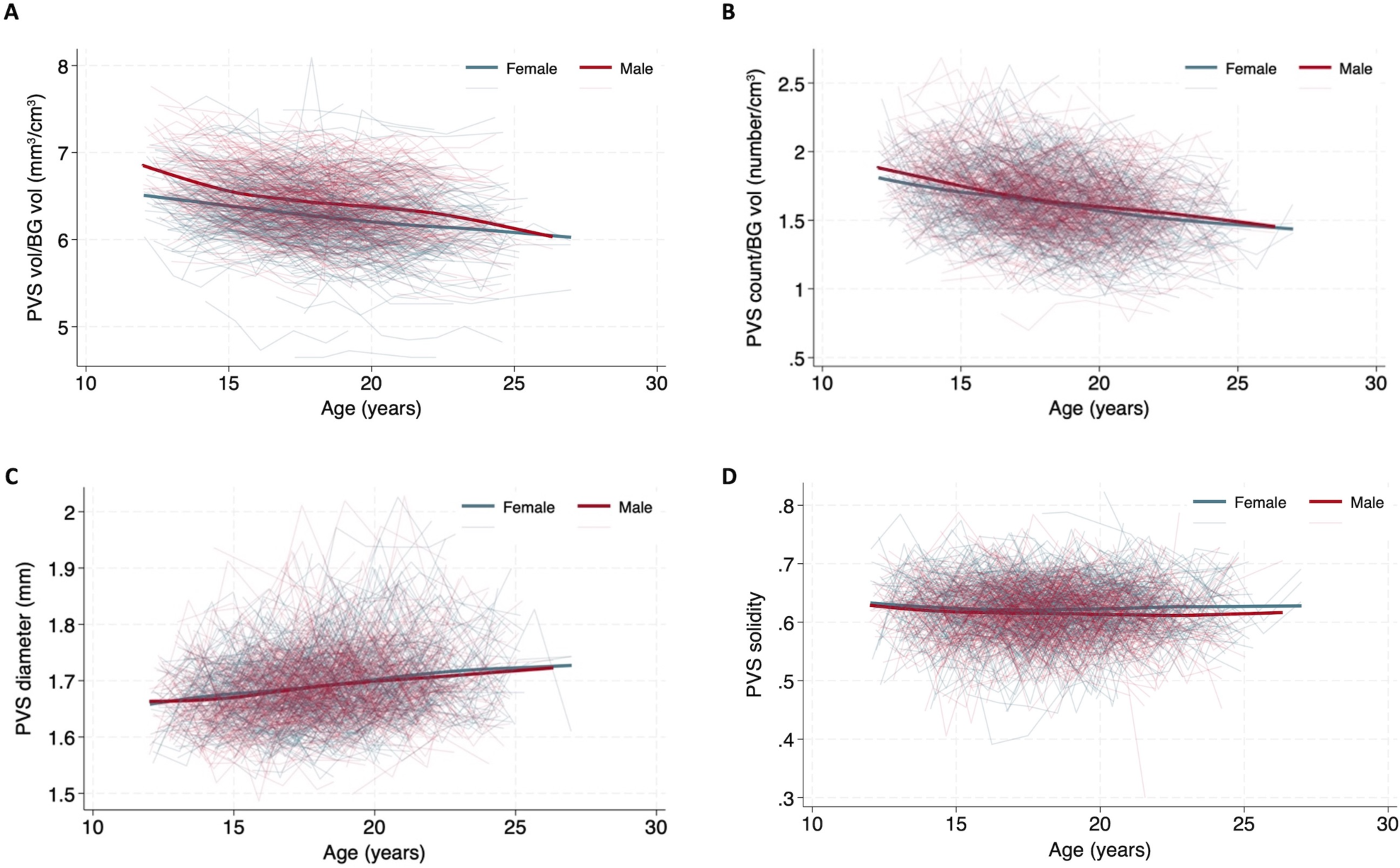
Relationship between basal ganglia MV-PVS burden and age. Changes in BG MV-PVS parameters A) volume B) count, C) diameter, D) solidity over time are represented for each individual by spaghetti plot. Locally weighted scatter plot smoothing curves derived from 3649 observations (1849 female-blue, 1800 male-red) are also shown. BG vol = basal ganglia volume.

### Modeling WM MV-PVS volume over time

Based on prior studies, MV-PVS volume was selected as the MV-PVS parameter of interest to evaluate the effects of time (age).[1,2] An unconditional random-intercept model of participant nested within site was first fit to the data. The intraclass correlation was 0.91, indicating that 91% of MV-PVS volume variance was due to person-mean differences. The remaining 10% of the variation was due to within-person residual variance as they age. Next, the effect of age on expected MV-PVS volume was analyzed via a sequence of mixed models. Inclusion of a fixed linear time coefficient was significant (*β*=0.13, SE=0.003, p<0.001). A random linear time slope variance (and covariance with the random intercept) significantly improved model fit, Δχ^2^(2)=202.4, p<0.001, indicating individual differences in linear rates of change. A fixed quadratic effect of age was also significant (*β*=-0.006, SE=0.001, p<0.001). Adding a random quadratic time slope variance (along with the two additional covariances, one with the random intercept and one with the random time slope) also improved model fit Δχ^2^(3)=19.5, p<0.001. The inclusion of higher-order fixed polynomial terms was not significant and the associated random effect term variances could not be estimated with adequate precision. Next, we assessed the effect of sex and BMI in the model. BMI was evaluated as two terms: person-mean BMI and person-occasion BMI.

An 18-year-old female was predicted to have a WM MV-PVS volume of 7.29 mm^3^/cm^3^, which was expected to increase by 0.156 mm^3^/cm^3^ each subsequent year (Supplemental Figure 3). Males showed a higher instantaneous rate of volume growth by age than females (0.183 mm/cm^3^/year versus 0.156 mm^3^/cm^3^/year, respectively, p<0.001). A quadratic main effect of time attenuated the instantaneous linear MV-PVS volume growth rate, indicating a deceleration in the rate of MV-PVS change over time. The MV-PVS volume growth rate of 0.16 mm^3^/cm^3^/year diminished by 0.004 per each additional year for females and by 0.011 for males (Supplemental Figure 3).

The fixed linear effect of age at 18 years old for males and females had 95% random linear age slope confidence intervals of (0.08, 0.28) and (0.05, 0.26), respectively, indicating the predicted range of the instantaneous effect of age on MV-PVS volume that 95% of the sample patients are predicted to fall into. The 95% random quadratic age confidence intervals for both sexes showed that while, on average, the rate of MV-PVS volume growth slows with time, for some subjects, the rate may increase.

### Association of BMI and sex on WM MV-PVS volume

The trajectory of WM MV-PVS volume over BMI was depicted by locally weighted scatter plot smoothing curves derived from individual subject observations by sex. In males, MV-PVS volume increased with BMI up to approximately 20 kg/m^2^, before reaching a plateau. In contrast, MV-PVS volume in females continued to increase with increasing BMI (Supplemental Figure 4)

The full linear mixed effects model for MV-PVS volume accounted for age, sex, person-mean BMI, person-occasion BMI, sex by age, sex by person-mean BMI, sex by person-occasion BMI, and scanner model (Supplemental Figure 4). Age, sex, person-mean BMI and person-occasion BMI were significantly associated with MV-PVS volume. We found statistically significant interactions of sex-by-linear age (*β*=0.028, SE=0.0075, p<0.001), sex-by-quadratic age (*β*=-0.0034, SE=0.0015, p=0.02), and sex-by-person-occasion BMI (*β*=-0.019, SE=0.006, p=0.001). Males aged 18 years, with both a person-mean BMI and person-occasion BMI of 23, displayed on average a 0.69 mm^3^/cm^3^ higher PVS volume burden than female counterparts of the same age and BMI (β=0.69, SE=0.076, p<0.001). However, in the female cohort, accounting for the effects of age, every one-unit increase in BMI above a person’s average BMI was associated with a 0.02 mm^3^/cm^3^ increase in MV-PVS volume (β=0.021, SE=0.005, p<0.001) (Figure 4). There was no significant relative effect of BMI in males.

**Figure 4.**
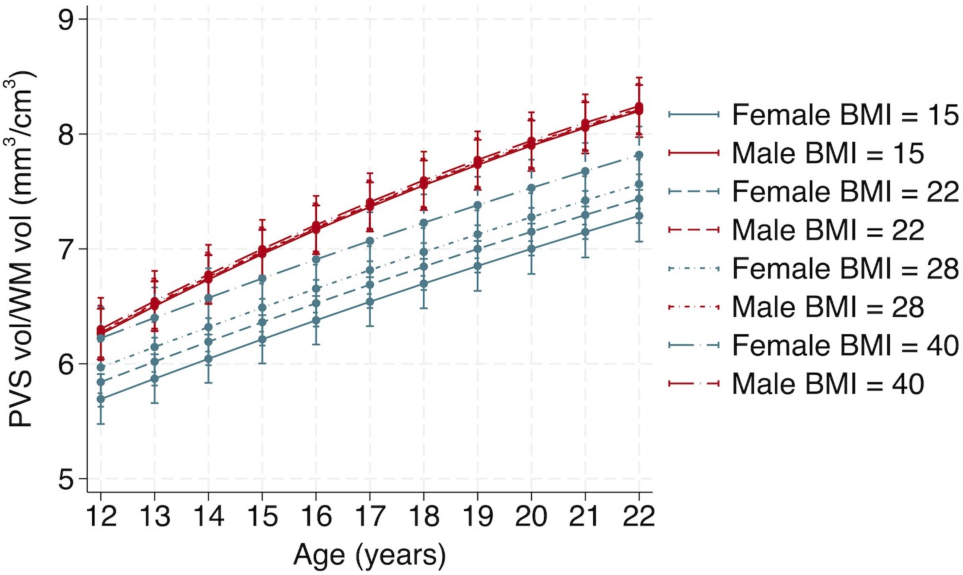
Relationship between MV-PVS volume and BMI. Line plots depicting the model predictions for male (red) and female (blue) MV-PVS volume by age with 95% confidence intervals. Predicted MV-PVS volume trajectories are replicated at 4 different person-occasion BMI values, which represent the effect of a deviation from an individual’s mean BMI on these predictions. BMI values were selected to reflect clinically meaningful categories of underweight (BMI 15), healthy (BMI 22), overweight (BMI 28), and obese (BMI 40) for an 18-year-old. WM vol = white matter volume.

## CONCLUSIONS

This is a robust longitudinal assessment of MV-PVS in adolescents and young adults. Our findings support prior cross-sectional analyses that MV-PVS burden increases with age,[1,2] young males have greater MV-PVS volume and count than females,[1,2] and that BMI is associated with MV-PVS volume.[28] Furthermore, we report that the MV-PVS volume growth rate slows quicker in males than females, and that increases in BMI were found to predict increases in MV-PVS volume in females only.

### The perivascular space as a marker of underlying biology

The perivascular space (PVS) is an anatomic compartment between penetrating blood vessels and brain parenchyma. In rodents, the PVS acts as a conduit for CSF flow between the subarachnoid space and the brain parenchyma.[7–9] Changes in MV-PVS volume, number, diameter, or solidity may reflect biological activity occurring in and around the PVS. The reason for PVS dilatation is unknown, but several theories exist, among them neuroinflammation, impaired CSF dynamics, and vascular pathology.[6]

### MV-PVS developmental trajectory

Despite growing evidence supporting the association between MV-PVS and neurologic pathologies in adults, MV-PVS are observed ubiquitously in the pediatric population. A previous large cross-sectional study across the lifespan (8-89 years) modeled a linear relationship between MV-PVS volume and age.[1] The current study is the first to provide longitudinal assessment of MV-PVS development over time, show the natural pattern of MV-PVS development, and to model changes in MV-PVS volume specifically within this demographic. We observed that while WM MV-PVS volume increases through adolescence, the rate of volume growth actually slowed during this specific period. With this data in hand, future studies may be able to determine an “average” or acceptable MV-PVS burden, and study MV-PVS burden in various pediatric pathologies relative to the healthy adolescent or young adult. Height and weight growth charts are essential components of pediatric care, and while not implemented clinically, expected developmental trajectory of the brain has also been mapped. Neither our study or the prior cross-sectional studies assess children younger than 8 years of age, and the prevalence and development of MV-PVS in the neonatal and young childhood ages remains understudied.[34]

There were striking differences between WM and BG MV-PVS developmental trajectories. Previous studies suggest that WM and BG MV-PVS may represent different underlying physiologies.[35] In our study, BG MV-PVS volume decreased with age. However, a previous study found that the relationship between BG MV-PVS volume and age was best described by a convex curve, with the nadir at 14 years, but this model included adult and elderly subjects who are anticipated to have higher BG MV-PVS burden.[1]

### Sex differences and MV-PVS

Our study highlights sex differences regarding: 1) total MV-PVS volume and number, 2) MV-PVS volume growth rate, 3) sensitivity of MV-PVS volume to changes in person-occasion BMI. Two cross-sectional studies of young subjects showed that males had greater MV-PVS volume and count than females.[1,2] Our analysis confirmed these differences and predicted males to have greater volume than females at all ages throughout the study. An analysis of 10 population-based adult cohort studies found that males had more WM MV-PVS until the 8^th^ decade of life when female MV-PVS count surged.[15] Changes in MV-PVS burden between males and females across the lifespan suggest that growth rate fluctuates over time. Indeed, we observed growth rate variability, with the rate of growth slowing more quickly in males than females. The role of sex-dependent biological factors in the progression of MV-PVSs has not been determined. In adolescence, the effect of puberty and associated hormonal changes should be investigated. Puberty in girls begins between 8 and 13 years,[36] just around study entry for the youngest participants (12 years). Of note, there was no statistical difference in MV-PVS volume between 12-year-old males and females, suggesting that this sex difference is not present earlier in childhood. Female puberty is associated with a rise in estrogen, which has affects systemic innate and adaptive immunity.[37] In the central nervous system, estrogen appears to exert a generally anti-inflammatory profile.[38] Thus sex hormones may influence inflammatory processes that are potentially relevant to the biology of enlarged PVS. However, sexual differentiation of the brain is likely more complex than linear downstream effects of sex hormones and has been proposed to involve multiple signaling pathways and epigenetic modifications.[39]

### MV-PVS and BMI

Our study demonstrates that in adolescence, higher mean BMI with age predicts greater MV-PVS volume over that same period. Furthermore, females are predicted to have additional increases in MV-PVS volume in response to rising BMI over time. The effect of BMI changes in females, although small, is consistent with a study reporting greater MV-PVS volume in females when BMI >30 kg/m^2^.[28] Additional studies should assess whether such small changes in MV-PVS volume holds clinical significance.

BMI is a clinical tool to categorize or diagnose obesity. Although changes in BMI are not synonymous with obesity, BMI may affect MV-PVSs through obesity-related biology. Obesity is a complex condition associated with adiposity, metabolic abnormalities, immune dysfunction, and chronic inflammation, which can affect brain development or physiology.[40,41] Obese adults (BMI 30-34 kg/m^2^) have higher inflammatory markers than nonobese counterparts, and females have greater inflammation than males in each BMI category.[42] Chronic inflammation in obesity is attributed to growing visceral adipocyte stores, that secrete proinflammatory cytokines and adipokines.[27] Inflammation can alter blood brain barrier function at the PVS through disruption of tight junctions, breakdown of the glia limitans, or changes in protein expression, which could cause accumulation of immune cells or fluid within the PVS.[43] Interestingly, obese female adolescents with visceral adiposity have relatively increased androgen activity,[44] which has been shown to promote CSF production.[45] Furthermore, rising BMI is associated with increasing CSF pressures, which could potentially result in dilation of CSF spaces such as the PVS.[26,46]

Based on our findings, we propose that each young individual sits upon a natural MV-PVS growth trajectory, and deviation from this trajectory may represent underlying biologic changes near the PVS that increase risk of additional “insult”. The relationship between female sex, BMI, and MV-PVS volume observed in this cohort is mirrored clinically by female-predominant neurologic conditions including idiopathic intracranial hypertension, neuroinflammatory disorders, and migraines. Idiopathic intracranial hypertension typically occurs in young females who are obese or with recent weight gain, and hormonal abnormalities (e.g. polycystic ovarian syndrome) are common comorbidities. In pediatric multiple sclerosis, obesity (defined as BMI ≥ 95 percentile by sex-specific BMI-for-age growth charts) was associated with an increased risk of multiple sclerosis or clinically isolated syndrome in girls, but not in boys.[47] The odds of episodic migraine is increased 81% in obese individuals compared to nonobese individuals, and this relationship was strongest in young women.[48] Each of these female-dominant neurologic conditions have been associated with increased MV-PVS burden.[22–25] The MV-PVS is a reflection of underlying subclinical changes in local biology likely associated with neuroinflammation, CSF and interstitial fluid exchange, or intracerebral solute clearance. The parallels between the female-dominant conditions above and their association with MV-PVS is interesting, and requires further study to determine the biological changes reflected by the MV-PVS, and whether such biology is related to disease pathophysiology in these disorders.

The greatest strengths of this study are its longitudinal design, ands the use of a large cohort of healthy adolescents and young adults which is generalizable to the greater youth community as the study cohort had a racial and ethnic composition comparable to the demographic of the United States.[29] However, this study has several limitations. This study cannot determine cause and effect and further experiments are needed to determine mechanistically how sex, BMI, and MV-PVS are linked. BMI was calculated from self-reported height and weight, which can introduce bias if inaccurately reported. Although we studied healthy participants, the variable BMI has a ceiling, in that participants with extremely high BMI may not fit in the MRI scanner and would be ineligible. Race, ethnicity, and socioeconomic status were not included as predictors in our model as there is conflicting evidence of any association with WM MV-PVS,[49,50] but should be considered in follow-up studies as these factors are known to influence regional brain growth.[51,52] Future studies must also evaluate how other clinical variables influence MV-PVS characteristics (e.g., blood pressure, sleep, headache, history of concussion, drug and alcohol use, and psychiatric comorbidities). Importantly, MV-PVS are small structures and the observed changes in MV-PVS volume produce small differences in slope. Although differences in slope of MV-PVS volume by age are significant, it remains to be determined whether these changes represent clinical significance.

## CONCLUSIONS

This longitudinal analysis demonstrates MV-PVS volume progression throughout adolescence and young adulthood, revealing sex and BMI differences in MV-PVS features. Future studies should assess whether puberty, sex hormones, and obesity play a role in this process, and whether deviation from one’s MV-PVS curve portends risk for conditions associated with increased MV-PVSs.

