## Supplemental for "Biological sex and BMI influence the longitudinal evolution of adolescent and young adult MRI-visible perivascular spaces"

### Supplemental Table1. Association between age, sex, BMI and MV-PVS volume over time

| | Estimate | Std. Error | 95% CI | | P-value |
|---|---|---|---|---|---|
| **Age** | 0.16 | 0.0055 | 0.14 | 0.17 | <0.001 |
| **Age by Age** | -0.0019 | 0.0010 | -0.0040 | -6e-06 | 0.05 |
| **Sex** | 0.69 | 0.076 | 0.54 | 0.84 | <0.001 |
| **Person-mean BMI** | 0.021 | 0.0089 | 0.0035 | 0.039 | 0.019 |
| **Person-occasion BMI** | 0.021 | 0.0042 | 0.013 | 0.029 | <0.001 |
| **Sex by Age** | 0.028 | 0.0075 | 0.013 | 0.042 | <0.001 |
| **Sex by Age by Age** | -0.0034 | 0.0015 | -0.0063 | -0.00058 | 0.02 |
| **Sex by occasion BMI** | -0.019 | 0.0060 | -0.031 | -0.0077 | 0.001 |
| **Scanner Model** | | | | | |
| **Prisma Fit** | -1.63 | 0.20 | -2.02 | -1.23 | <0.001 |
| **SIGNA Creator** | -0.16 | 0.66 | -0.19 | -0.037 | 0.01 |
| **Trio Tim** | -1.05 | 0.20 | -1.43 | -0.66 | <0.001 |

Age is centered around 18 years. Person-occasion BMI is centered at 23 kg/$m^2$.

**Supplemental Figure 1. Consort flow diagram of participant eligibility.**

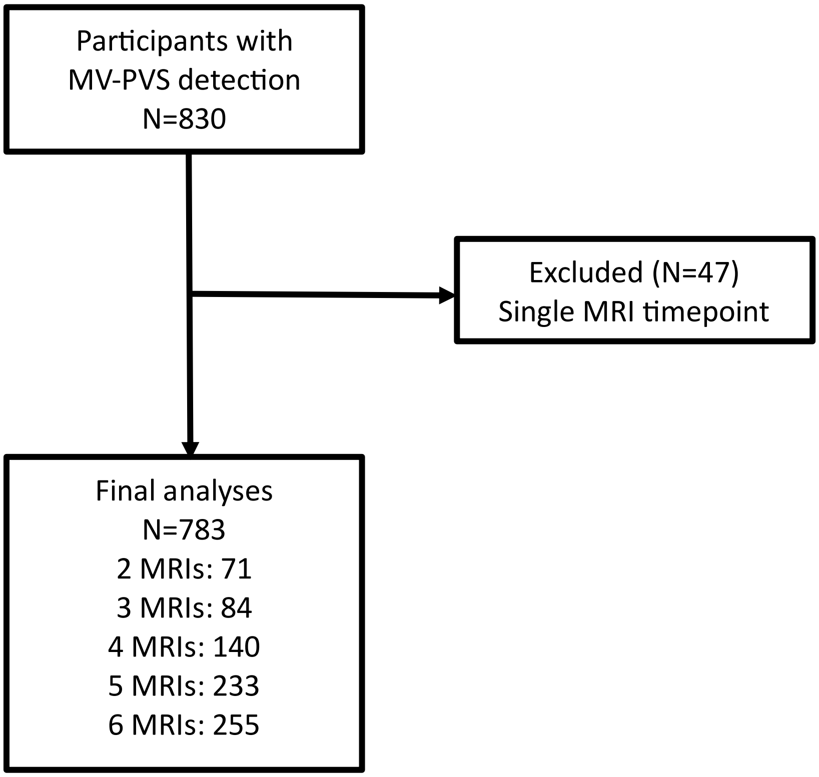

**Supplemental Figure 2. White matter volume increases with age in both males and females.** Changes in supratentorial white matter volume over time are represented for each individual by spaghetti plot. LOWESS curves derived from 3649 observations (1849 female-blue, 1800 male-red) are also shown.

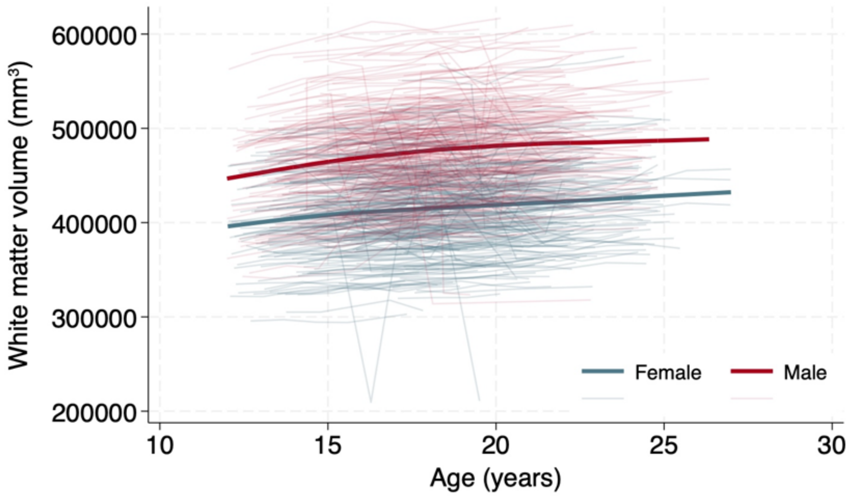

**Supplemental Figure 3. Linear mixed model predictions of MV-PVS volume.** A) Expected MV-PVS volume over time for an individual, accounting for BMI. B) Instantaneous linear rate of expected MV-PVS volume growth declines over time. Shaded region marks 95% confidence interval (blue- female, red- male).

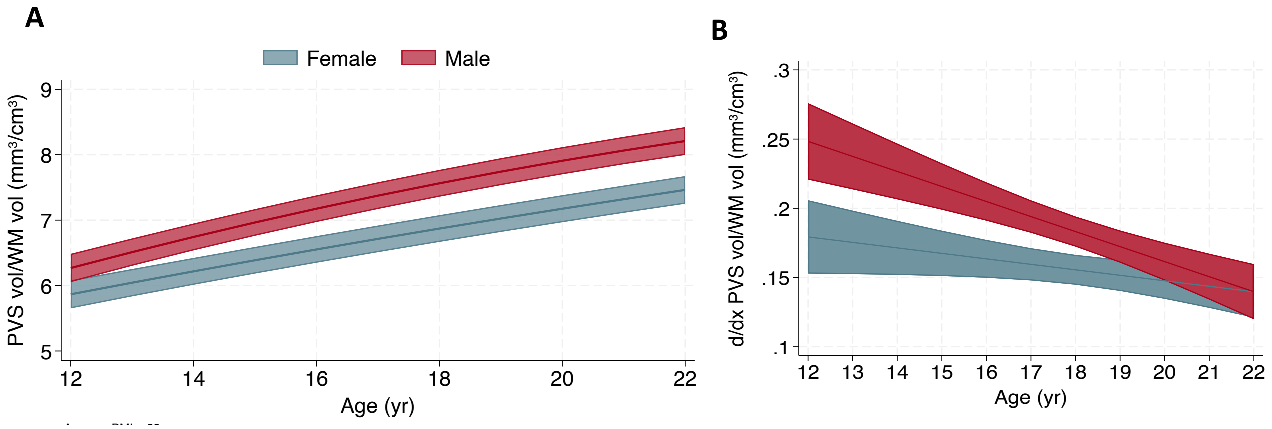

**Supplemental Figure 4. Relationship between BMI and MV-PVS volume over time.** Changes in WM MV-PVS volume by BMI over time are represented for each individual by spaghetti plot. LOWESS curves derived from 3649 observations (1849 female-blue, 1800 male-red) are also shown. WM vol = white matter volume.

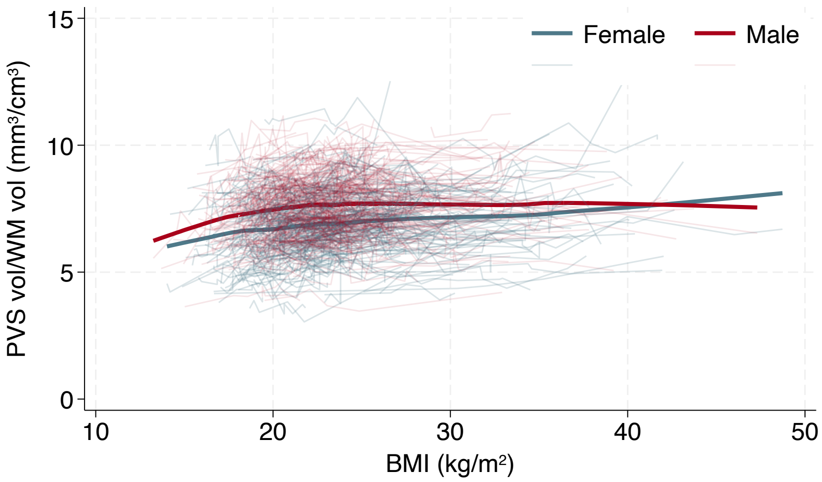
